# Inhaled nicotine equivalent to cigarette smoking disrupts systemic and uterine hemodynamics and induces cardiac arrhythmia in pregnant rats

**DOI:** 10.1101/195099

**Authors:** Xuesi M. Shao, Héctor E. López-Valdés, Jing Liang, Jack L. Feldman

## Abstract

Maternal smoking with obligatory nicotine inhalation is associated with preterm delivery, low birth weight, fetal growth retardation and developmental defects. We tested the hypothesis that cigarette smoking-relevant nicotine inhalation during pregnancy impairs cardiovascular function and uterine hemodynamics with consequential fetal ischemia. Pregnant rats exposed to episodic inhaled nicotine via a novel lung alveolar region-targeted aerosol method produced nicotine pharmacokinetics resembling cigarette smoking in humans. This clinically relevant nicotine aerosol inhalation (NAI) induced transient reduction and irregular fluctuations in uterine artery blood flow associated with cardiac arrhythmia and high magnitude irregular fluctuations of systemic blood pressure. The arrhythmia included sinoatrial (SA) block, sinus arrest, 2^o^ and 3^o^ atrioventricular (A-V) block and supraventricular escape rhythm. These effects were blocked by the nicotinic receptor (nAChR) antagonist mecamylamine. Resection of the ovarian nerve, which innervates uterine blood vessels, counteracted the NAI-induced reduction in uterine blood flow. We suggest that the rapid rise pattern of arterial blood nicotine concentration stimulates and then desensitizes autonomic nAChRs leading to disruptions of cardiac function as well as systemic and uterine hemodynamics that reduces uteroplacental blood flow, a mechanism underlying maternal smoking-associated pregnancy complications and developmental disorders. These findings challenge the safety of pure nicotine inhalation, i.e., E-cigarettes.

## Introduction

Maternal smoking increases the risk of preterm delivery, low birth weight, fetal growth retardation, sudden infant death syndrome (SIDS) and developmental defects ^1^. The mechanisms are not clear. Smoking induces acute cardiovascular events including tachycardia, high arterial blood pressure (BP), coronary spasm, and increases in myocardial work and oxygen demand with concomitant reduction in blood and oxygen supply ^2^. Cigarette smoking is associated with sudden cardiac deaths, most of which are due to arrhythmias ^2,3^. Smoking cessation reduces cardiac deaths from arrhythmia for patients with cardiac conditions ^4^. Nicotine, the major bioactive component of cigarette smoke, is the most significant contributor to these acute cardiovascular effects ^5^. With the recent emergence and increasing popularity of devices for inhalation of nicotine, e.g., non-combustion inhaler type of tobacco products ^6^ and electronic nicotine delivery system/e-cigarettes, often portrayed as safer alternatives to cigarettes and cigars, understanding the effects of inhaled nicotine faces an increasing urgency.

One requirement for animal studies of drugs of abuse to be relevant to humans is that the route of drug administration as well as the blood pharmacokinetics (PK) or target tissue concentrations are comparable ^7,8^. In previous mammalian studies on the role of nicotine in the adverse effects of tobacco smoking on circulation, including on uterine circulation, and fetal development, there were major limitations: i) routes of nicotine delivery, e.g., oral application, intraperitoneal (i.p.), intravenous (i.v.) injections, or subcutaneous osmotic pump, resulted in nicotine blood PK distinctly different from human smoking, including the rapidity of absorption, rise time and peak of arterial blood concentrations, which determine the magnitude of some cardiovascular effects of nicotine ^9^. ii) Blood flow and arterial BP measurements were not continuous, so that transient events could be missed. iii) The effects on uterine blood flow have been inconclusive, due to the indirect nature of the measurements, i.e., flow was calculated from vessel diameter measured with ultrasound and blood velocity measured with Doppler velocimeter. However, since flow is proportional to the fourth power of vessel diameter, even small errors in the diameter measurement would cause large deviations in the estimate of blood flow. Alternative parameters such as the ratio of systolic to diastolic blood flow velocity (S/D ratio), pulsatile index and resistance index of uterine artery are all based on velocity, not volume flow that directly reflects uterine blood supply ^10^. Here, we delivered nicotine to pregnant rats with the lung alveolar region-targeted aerosol technology ^11^ while continuously monitoring electrocardiogram (ECG), systemic BP along with making high-resolution direct measurements of uterine artery blood flow ^12^. We tested the hypothesis that nicotine NAI exposure in dose and kinetics equivalent to that in human smoking stimulates autonomic nAChRs resulting in disturbances in cardiac function and systemic hemodynamics as well as vasoconstriction of the uterine artery that disrupts uteroplacetal hemodynamics, which can lead to fetal ischemia. This provides a mechanistic link between maternal smoking and its adverse effects on pregnancy and fetal development.

## Results

### Nicotine aerosol inhalation produces pharmacokinetics in pregnant rats resembling that of human cigarette smoking

Nicotine aerosol in the breathing zone of the nose-only exposure chamber in our aerosol system has a droplet size distribution within the respirable diameter range ^13,14^ that results in deposition of nicotine in lung alveolar regions and rapid absorption into the pulmonary circulation, and then rapidly into the coronary and systemic circulations ^11^. Pregnant rats were exposed to NAI for 2 min. The arterial blood nicotine concentration reached a maximum (C_max_) of 35 ± 12.3 ng/ml (mean ± SD) within 1–3 min (T_max_ = 1 min), declined quickly to 14 ± 2.6 ng/ml and then ranged to 12-15 ng/ml for up to 40 min (Fig. 1 A). Venous blood plasma nicotine levels gradually increased to 14 ± 4.4 at 25 min, close to the arterial nicotine level (Fig. 1 A). Arterial and venous levels of cotinine (the predominant metabolite of nicotine) increased slowly to 11–16 ng/ml at 40 min (Fig. 1 B). The concentrations and temporal profile (PK) of nicotine in arterial and venous blood in NAI exposure rats closely resembled those of humans smoking a cigarette ^15,16^.

**Figure 1.**
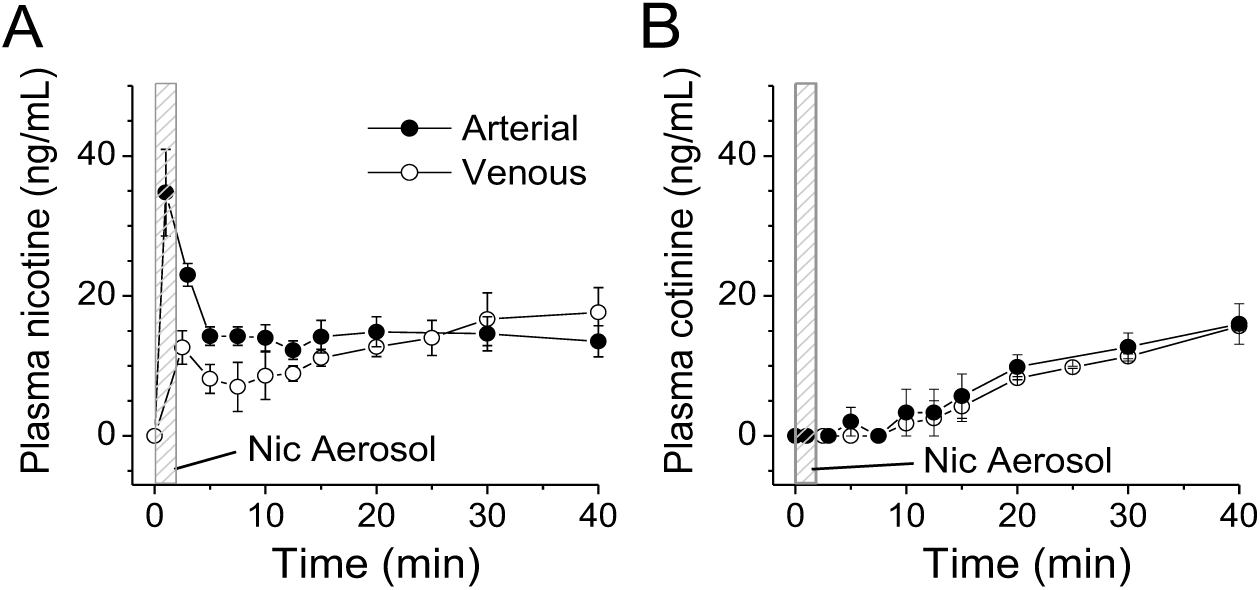
Nicotine aerosol inhalation (NAI) exposure in pregnant rats produces blood PK in arterial and venous blood resembling humans smoking a cigarette^15^. (**A**) Plasma nicotine levels (mean ± SE, n = 4). (**B**) Plasma cotinine (a major nicotine metabolite) levels. Rats were awake, constrained in a rat holder and expose to aerosol with a nose-only chamber for 2 min (indicated as a gray area in (A) and (B)). Nicotine concentration in the nebulizer was 1%. Blood samples were collected from the femoral artery or the tail vein (separate rats) in a series time points from the start of nicotine aerosol delivery (time 0) including one sample prior to the start of NAI (the data is shown at time point 0).

### NAI induces a hemodynamic disruption in uterine circulation correlated with cardiac arrhythmia

To understand how episodic inhaled nicotine, in PK equivalent to that of human smoking, affects the uterine blood supply in pregnant rats, we measured uterine artery blood flow with a chronically implanted perivascular micro-flowprobe while recording ECG. Pre-aerosol mean blood flow was 1.52 ± 0.69 ml/min, with a regular pulsing of 7.0 ± 0.86 Hz synchronized with ECG waves (Fig. 2 A, D and E). The diastolic flow was 0.97 ± 0.47 ml/min and pulse flow 1.57 ± 0.56 ml/min (Fig. 2 F and H). NAI exposure reduced mean uterine blood flow to 0.82 ± 0.61 ml/min (50 ± 21% of pre-nicotine control) and pulsing frequency to 4.1 ± 0.64 Hz (Fig. 2 B, D and E). Diastolic flow was reduced to 0.58 ± 0.47 ml/min and the pulse flow reduced to 0.91 ± 0.64 ml/min (Fig. 2 F and H). Fluctuations in diastolic and pulse flow were substantial (Fig. 2 B), as quantified by the coefficient of variation (CV). CV of diastolic flow increased from 0.098 ± 0.032 to 0.44 ± 0.25 (pre-nicotine vs. NAI, 4.5-fold increase) (Fig. 2 G). The irregularity of pulse amplitude also greatly increased as its CV changed from 0.064 ± 0.021 to 0.38 ± 0.21 (5.9-fold increase, Fig. 2 I).

**Figure 2.**
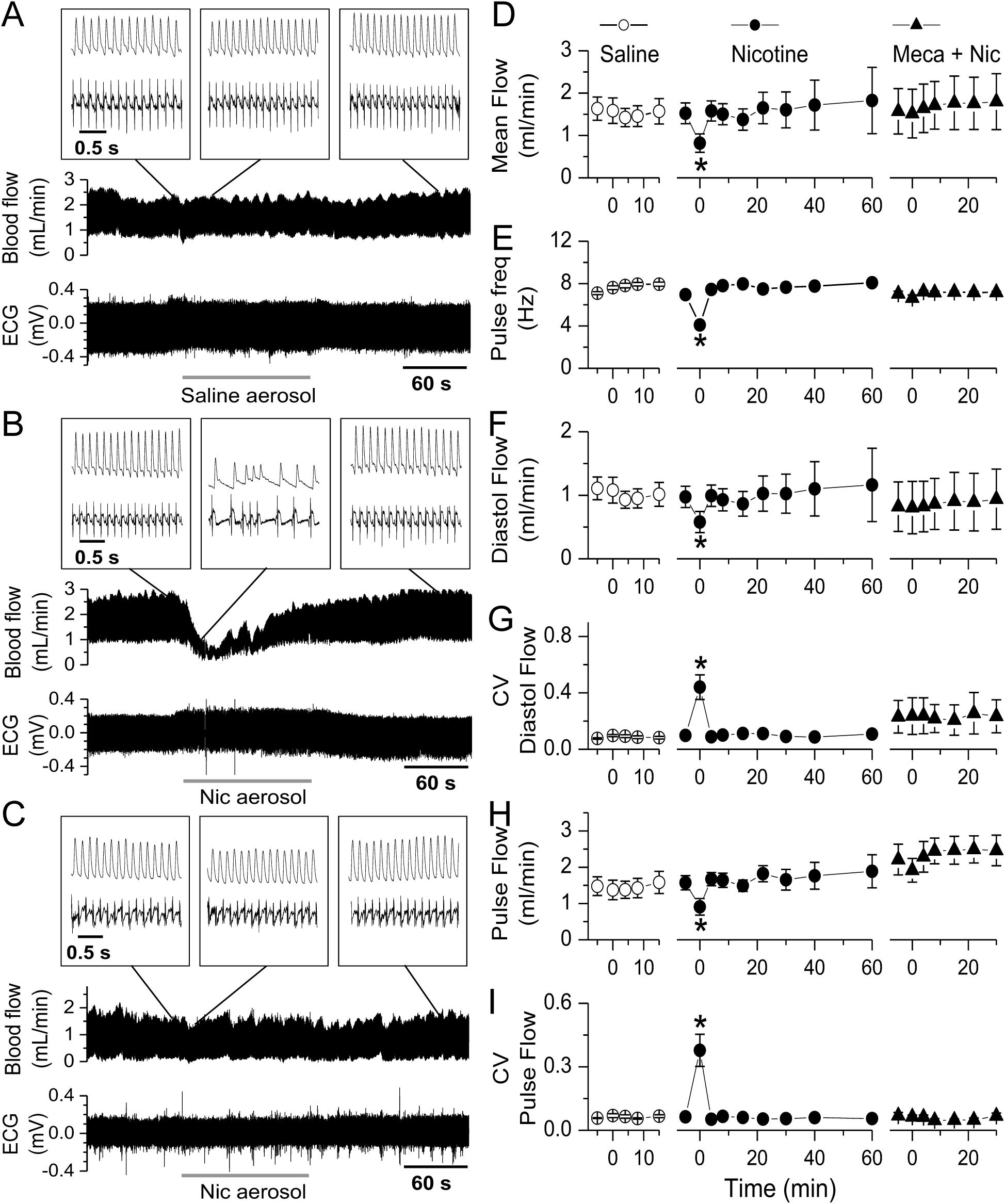
NAI in pregnant rats with a PK equivalent to smoking a cigarette in humans induces a hemodynamic disruption in the uterine artery and cardiac arrhythmia. Simultaneous recordings of ECG and uterine artery blood flow from rats. Aerosol was generated as described in Materials and Methods. (**A**) Saline aerosol inhalation (2 min). (**B**) Nicotine (Nic) aerosol exposure induced a transient reduction in blood flow and pulsing frequency, and induced fluctuations in diastolic flow and irregular pulse flow in the uterine artery as well as induced cardiac arrhythmia. The arterial blood flow pulse was still synchronized with ECG waves. (**C**) These NAI effects were eliminated in the presence of a nAChR antagonist mecamylamine (Meca, 3mg/kg, i.p. injection). Insets are sections of traces with expanded time scale from the main traces at time points indicated by straight lines. Summary parameters for the effects of aerosol inhalation/exposure on uterine artery blood flow including saline aerosol (n = 7), Nic aerosol (n = 8) and Nic aerosol in the presence of Meca (Meca + Nic, n = 3): (**D**) Mean blood flow. (**E**) Flow pulse frequency (freq). (**F**) Diastolic flow (minimum before rising of each pulse). Coefficient of variation (CV) of diastolic flow. (**H**) Pulse flow (the amplitude differences between diastolic flow and maximum of each pulse). (**I**) CV of pulse flow. Time 0 is the onset time of aerosol exposure. During aerosol exposure of 2 min, the data point is a sample of 30 sec starting 5 to 10 sec after the onset of aerosol delivery. *, statistical significance (p<0.05) of the parameter during NAI vs. pre-nicotine conditions.

ECG QRS waves were synchronized, with a time delay, with blood flow pulsing. When the ECG interpulse, i.e., R-R, intervals, lengthened, blood flow pulsing frequency and blood flow dropped; when the heart rate increased, the blood flow pulsing frequency and flow increased (Fig. 2 B middle inset).

These effects were transient, starting at 4.4 ± 2.8 sec from the onset of NAI, lasting 78.2 ± 10.7 sec in the presence of nicotine aerosol (Fig. 2 B).

The regularity of blood flow was quantified with time-domain analysis. The normalized autocovariance function (NACVF, equation (2) in Methods) showed periodic flow in pre-nicotine conditions (Fig. 3 A). During NAI, the autocorrelation dropped quickly with time lag, and the periodic correlation diminished (Fig. 3 B). The power spectrum showed a frequency component of ∼6-8 Hz in pre-nicotine conditions that diminished during NAI (Fig. 3 A and B), suggesting a loss of periodicity in uterine blood flow.

**Figure 3.**
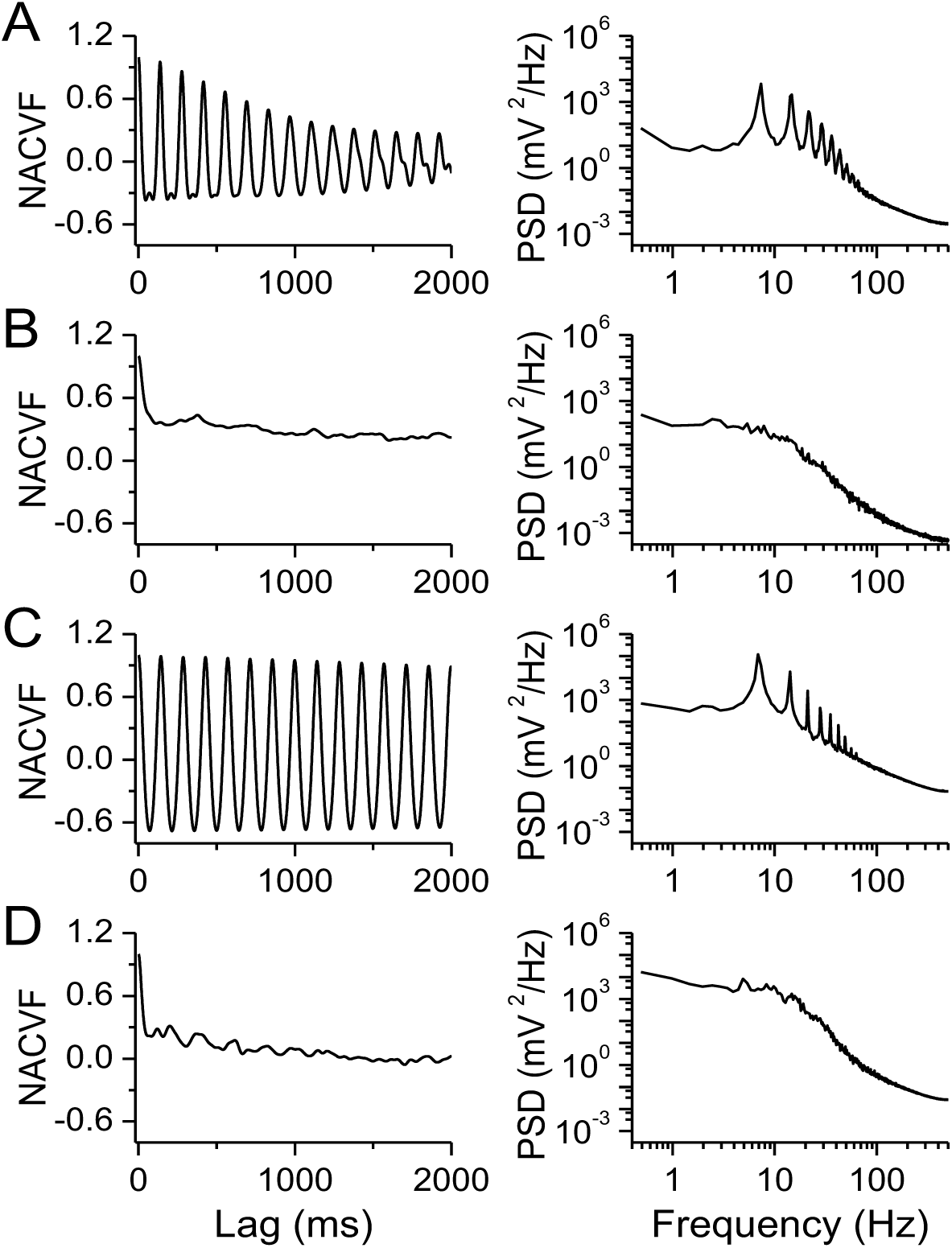
NAI exposure diminishes periodicity of uterine artery blood flow. Mecamylamine (Meca) administration counteracts this effect while resection of the ovarian nerve does not affect the nicotine effect on periodicity. Left panels are normalized autocovariance function (NACVF) and right panels are power spectral density (PSD) of blood flow. (**A**) Pre-nicotine conditions. (**B**) NAI exposure. (**C**) NAI in the presence of Meca (3mg/kg, i.p. injection). (**D**) NAI with the ovarian nerve cut.

To quantify the temporal relationship between uterine artery blood flow and cardiac activity, we calculated the normalized cross-covariance function (NCCVF, equation (4) in Methods) of blood flow vs. ECG (Fig. 4). NCCVF was high and periodic in pre-nicotine conditions, consistent with blood flow being highly correlated with the rhythmic heartbeat. During NAI, NCCVF had a high cross-correlation within one cycle then rapidly diminished (Fig. 4 A and B). Blood flow periodicity was disturbed while the cross-correlation with cardiac activity persisted. These results suggest that NAI-induced cardiac arrhythmia and irregular cardiac output disrupted rhythmic uterine blood flow and hemodynamics.

**Figure 4.**
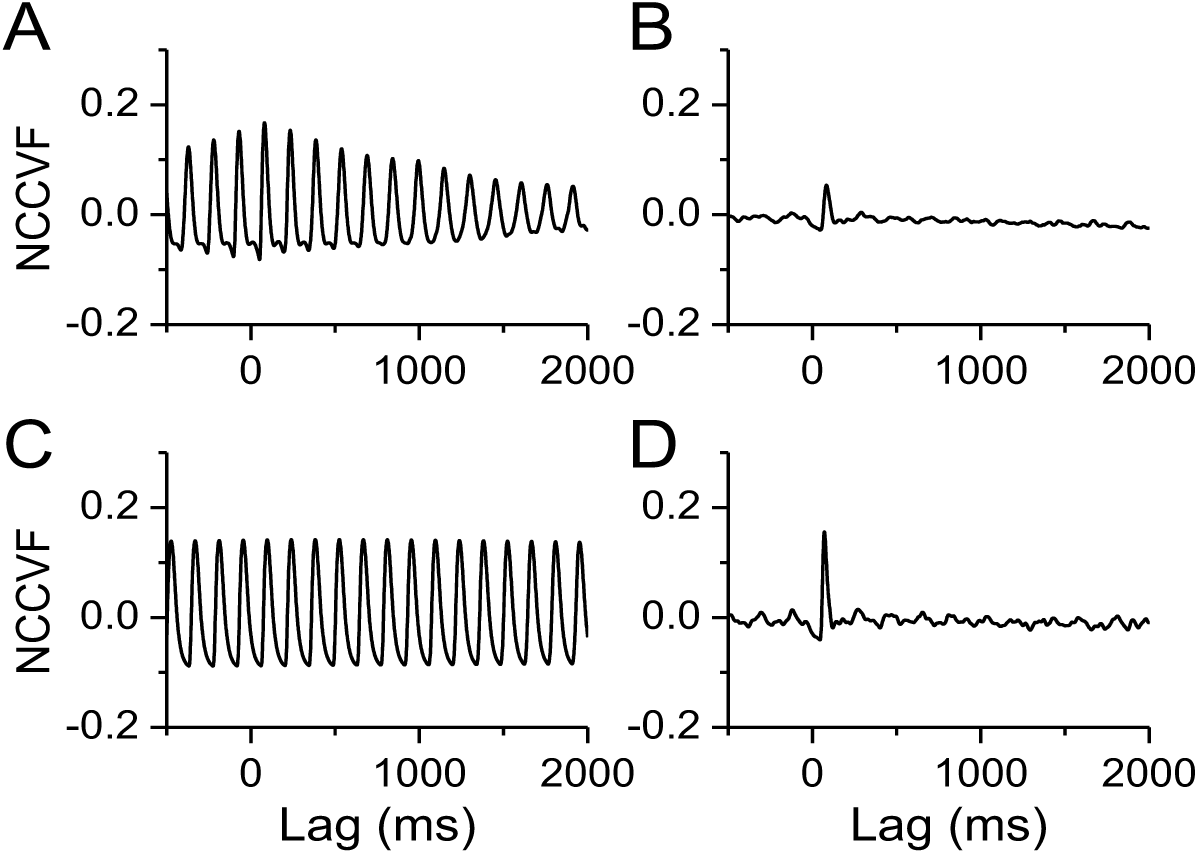
NAI diminishes periodicity of uterine blood flow while cross-correlation remains within one cycle. Meca blocks the NAI effects while NAI effect on cross-correlation persists following resection of the ovarian nerve. Normalized cross-covariance function (NCCVF) of uterine blood flow as a continuous process vs. ECG as a point process was calculated. (**A**) NCCVF in pre-nicotine conditions. (**B**) NAI exposure. (**C**) NAI in the presence of Meca. (**D**) NAI with the ovarian nerve cut.

Following systemic administration of the nAChR antagonist mecamylamine (Meca), NAI had no effects on mean blood flow, pulse frequency, diastolic or pulse flow, nor on their rhythmicity (Fig. 2 C-I); Meca also eliminated the effects of NAI on heart rate and the temporal relationship between uterine blood flow and heartbeat (Fig. 3 C and Fig. 4 C).

These results demonstrate that clinically relevant NAI exposure in pregnant rats stimulates autonomic nervous system nAChRs that modulate cardiac function and vascular tone, inducing transient bradycardia, arrhythmia, uterine artery vasoconstriction and, as a consequence, local hemodynamic disruption of uterine circulation including reduction, fluctuation and irregular pulsing of uterine artery blood flow.

### Resection of the ovarian nerve counteracts NAI-induced reduction, but not irregular fluctuation, of uterine artery blood flow

To investigate the role of neural control in mediating the effects of nicotine on uterine blood flow, we unilaterally resected the ovarian nerve that innervates uterine arteries. NAI reduced mean blood flow from 1.08 ± 0.34 to 0.68 ± 0.26 mL/min, i.e., a reduction to 63 ± 11% with the nerve cut compared to a reduction to 50 ± 21% with the nerve intact (p-value of the contrast of the difference between pre- and during NAI with nerve cut vs. the difference between pre- and during NAI with nerve intact is 0.0068 in a two-way repeated measures ANOVA model), and reduced NAI effects on diastolic flow and variability of pulse flow of the uterine artery (Fig. 5 A, B, C and F vs. Fig. 2 B, D, F and I). Resecting the ovarian nerve had no statistically significant effect on: NAI-induced decrease in pulse flow; increase in variability of diastolic flow (Fig. 5 E and D, and Fig. 2 H and G); periodicity of uterine artery blood flow, and; crosscorrelation between heartbeat and uterine artery blood flow (Fig. 3 D and Fig. 4 D). These results indicate that NAI-induced reduction of uterine artery blood flow is partially mediated by ovarian nerve from excitation of sympathetic nAChRs^17^. The auto- and cross-correlation analysis further suggests that the NAI-induced irregular fluctuation of blood flow is primarily due to cardiac arrhythmia which is not affected by the ovarian nerve.

**Figure 5.**
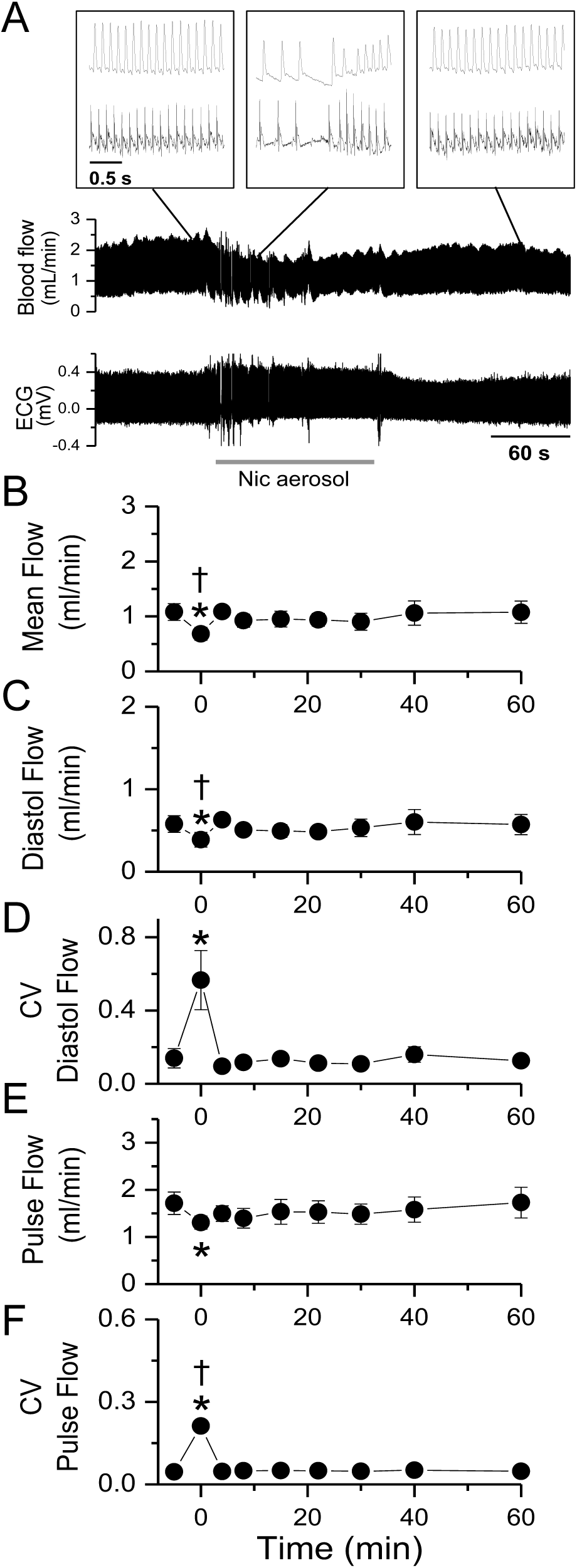
Unilateral section of the ovarian nerve counteracted the effects of NAI on uterine artery blood flow in pregnant rats ECG was simultaneously recorded. (**A**) Nicotine (Nic) aerosol exposure induced a transient reduction in blood flow and pulsing frequency, fluctuations in diastolic flow and irregular pulse flow in the uterine artery as well as cardiac arrhythmia. Summary parameters: (**B**) mean blood flow. (**C**) Diastolic (Diastol) flow. (**D**) CV of diastolic flow. (**E**) Pulse flow. (**F**) CV of pulse flow. *, statistical significance (p<0.05) of the parameter during NAI vs. pre-nicotine conditions (n = 5). †, statistically significant contrast of difference between pre- and during NAI with ovarian nerve section vs. difference between pre- and during NAI with ovarian nerve intact (two-way repeated measure ANOVA followed by post hoc multiple comparison analysis Sidak method).

### NAI induces a high magnitude irregular fluctuation of systemic BP correlated with cardiac arrhythmia

To understand how NAI disturbs systemic hemodynamics, we continuously measured arterial BP. In pre-nicotine control conditions, mean BP was 105.5 ± 11.3 mmHg, and mean diastolic pressure, 102.2 ± 11.4 mmHg with a pulse pressure of 5.4 ± 2.7 mmHg and pulse frequency of 8.0 ± 0.54 Hz that was synchronized with ECG waves (Fig. 6 A and D). From the onset of NAI exposure, there were progressive, large fluctuations of BP and obvious ECG rhythm changes. When the heartbeat was fast, BP increased substantially (Fig. 6 B), and when it was slow, BP dropped sharply. The nicotine-induced high magnitude fluctuations of BP were reflected in the CV of diastolic pressure (0.14 ± 0.039 vs. 0.022 ± 0.005 in pre-nicotine control, Fig. 6 G) and CV of pulse pressure (0.44 ± 0.17 (vs. 0.097 ± 0.035 in pre-nicotine control, Fig. 6 I). With continued NAI, BP fluctuations subsided at a slightly higher BP level that extended to ∼46 ± 16 sec after nicotine exposure. NAI also induced cardiac arrhythmia (Fig. 6 B), a decrease in pulse BP frequency to 4.1 ± 0.39 Hz (Fig. 6 E) and an increase in pulse pressure to 9.6 ± 4.5 mmHg (Fig. 6 H). The effect was transient, starting 3.2 ± 0.9 seconds from NAI onset and lasting 126 ± 11 s. NAI effects on mean BP and diastolic BP during this epoch were not statistically significant (Fig. 6 B, D and F).

**Figure 6.**
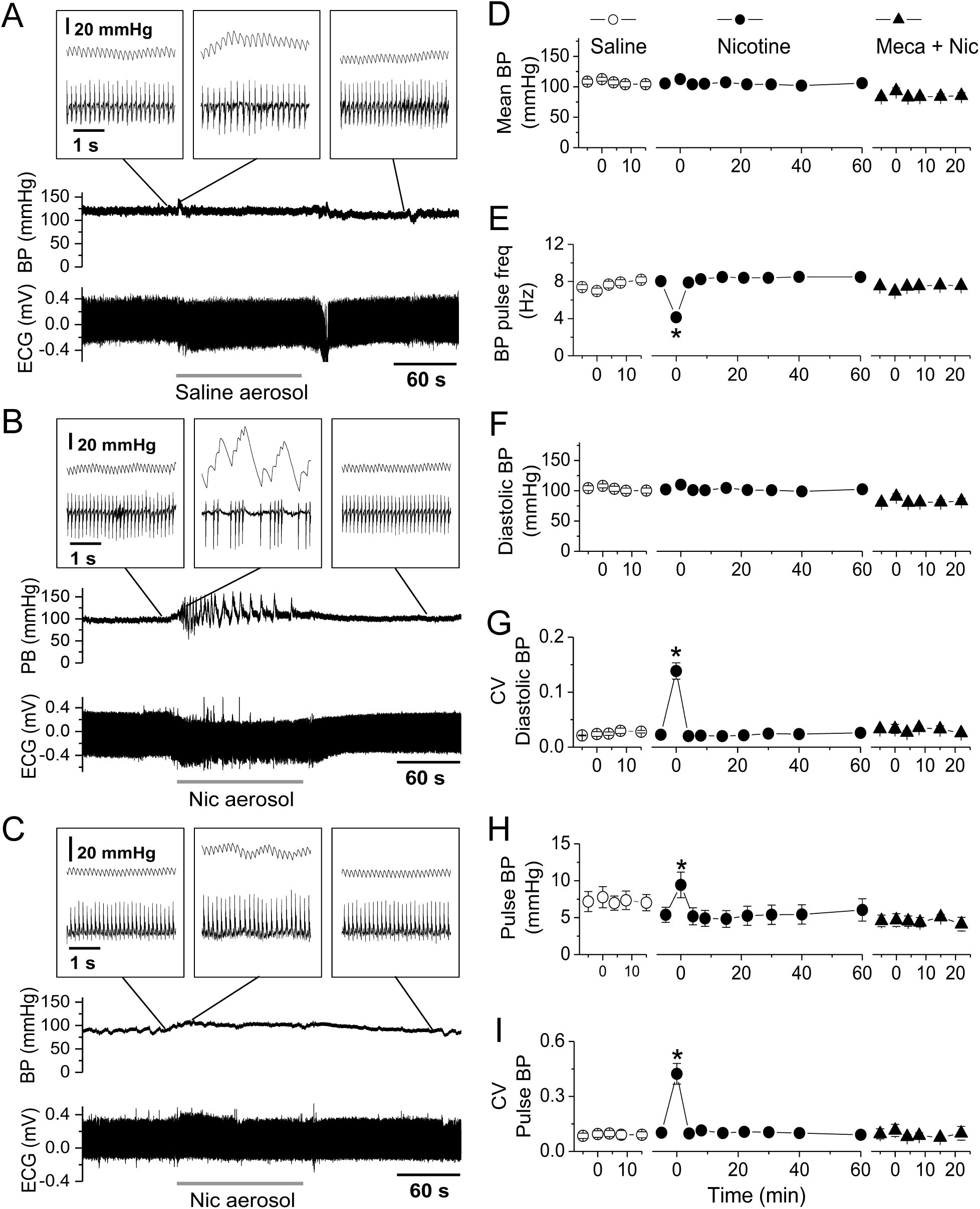
NAI in pregnant rats with a PK equivalent to smoking a cigarette in humans induces a high magnitude fluctuation of BP and cardiac arrhythmia. Simultaneous recordings of ECG and BP from rats. (**A**) Saline aerosol inhalation. (**B**) Nicotine (Nic) aerosol exposure induced a high magnitude fluctuation in diastolic BP, reduction in BP pulsing frequency and irregular pulse BP as well as induced cardiac arrhythmia. Note that the PB pulse was synchronized with ECG waves. (**C**) Effects of NAI on BP and ECG were eliminated in the presence of Meca (3mg/kg, i.p. injection). Insets are sections of traces with expanded time scale from the main traces at time points indicated by straight lines. Summary parameters (mean ± SE) for the effects of aerosol on BP including saline aerosol (n = 5), nicotine (Nic) aerosol (n = 7), Nic aerosol in the presence of Meca (Meca + Nic, n = 3): (**D**) Mean BP. (**E**) BP pulsing frequency. (**F**) Diastolic pressure. (**G**) Coefficient of variation (CV) of diastolic pressure. Pulse pressure. (**I**) CV of pulse pressure. *, statistical significance (p<0.05) of the parameter during NAI vs. pre-nicotine conditions.

As cholinergic transmission mediated by nAChRs is fundamental for BP regulation, systemic Meca decreased mean BP and diastolic BP (Fig. 6 D and F) while eliminating the effects of NAI on pulse frequency, high magnitude BP fluctuations, pulse pressure, CV of diastolic BP and CV of pulse pressure. These data indicate that the effects of clinically relevant inhaled nicotine on cardiac function, hemodynamic and BP are mediated by nAChRs.

NAI exposure induced (based on ECG morphological analysis; Fig. 7): i) slow and irregular sinus p-waves, deformed ectopic p-waves, some of which were non-conducted, sinoatrial (SA) block; ii) atrioventricular (AV) junctional or atrial escape rhythm with 2^o^ AV block (Fig. 7 B); iii) SA block and sinus arrest (Fig. 7 C); iv) paroxysmal supraventricular tachycardia (SVT) with AV block (Fig. 7 D); v) independent series of p-waves and QRS complexes suggesting 3^o^ AV block and AV junctional escape rhythm (Fig. 7 E), and; vi) ventricular premature beats (Fig. 7 F) in pregnant rats. Following administration of Meca, regular sinus rhythm persisted upon NAI (Fig. 7 G).

**Figure 7.**
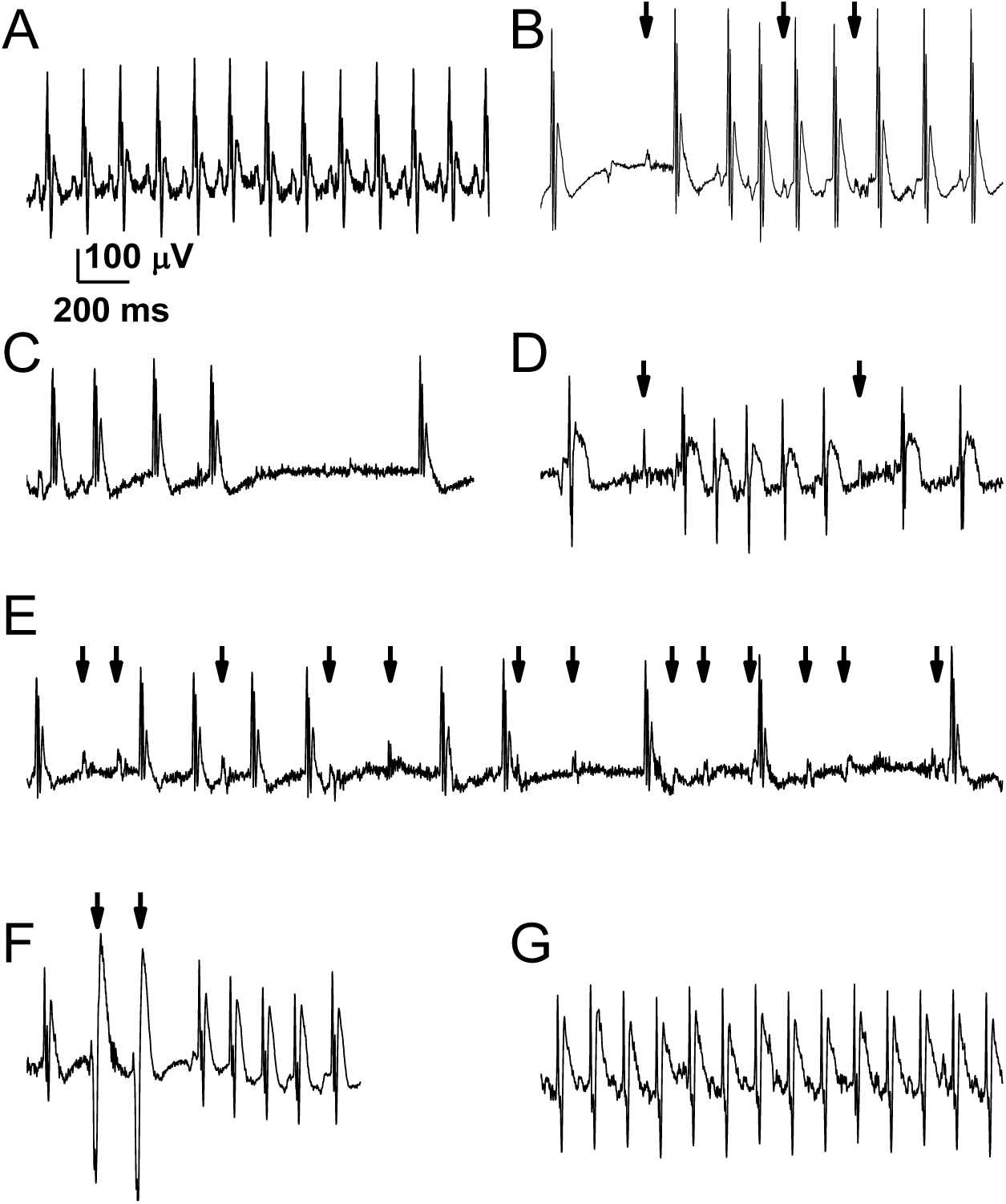
NAI effects on ECG morphology and rhythm (lead III). (**A**) Regular sinus rhythm in control (pre-nicotine) conditions. During NAI: (**B**) Sinoatrial (SA) block and atrioventricular (AV) junctional or atrial escape rhythm with AV block 2^o^. Arrows indicate sinus P waves which are slow and irregular of which the 1^st^ and 3^rd^ are non-conducted, each followed by an AV junctional escape beat. Other P waves were multifocal atrial waves that contributed to the escape rhythm; (**C**) SA block and sinus arrest. Two sinus beats, then SA block plus an AV junctional escape beat, a sinus beat followed by a sinus arrest lasting 812 ms, then an AV junctional escape beat; (**D**) Paroxysmal supraventricular tachycardia (SVT) with AV block. An atrial ectopic beat, then a non-conducted sinus P wave (arrow) followed by an atrial ectopic beat and a SVT. Then a 2^nd^ non-conducted sinus P wave (arrow) followed by two AV junctional escape beats; (**E**) AV block 3^o^ and AV junctional escape rhythm. P waves (arrows) are independent of QRS complexes; (**F**) Ventricular premature beats (VPBs). An AV junctional escape beat, 2 VPBs (arrows) followed by a SVT. (**G**) Regular sinus rhythm persists when the rat exposure to nicotine in the presence of mecamylamine. x-y scale is the same for all panels. n = 8 rats.

We suggest that clinically relevant NAI-induced bradycardia and arrhythmia are primarily due to SA block and AV block while dominated by supraventricular escape rhythm, signs consistent with elevated vagal tone.

During NAI exposure, BP autocorrelation dropped with time lag, and periodic correlation diminished (Fig. 8A and B). The power spectrum peaked ∼6-8 Hz in pre-nicotine conditions diminished during NAI (Fig. 8 A and B), indicating a loss of BP periodicity.

**Figure 8.**
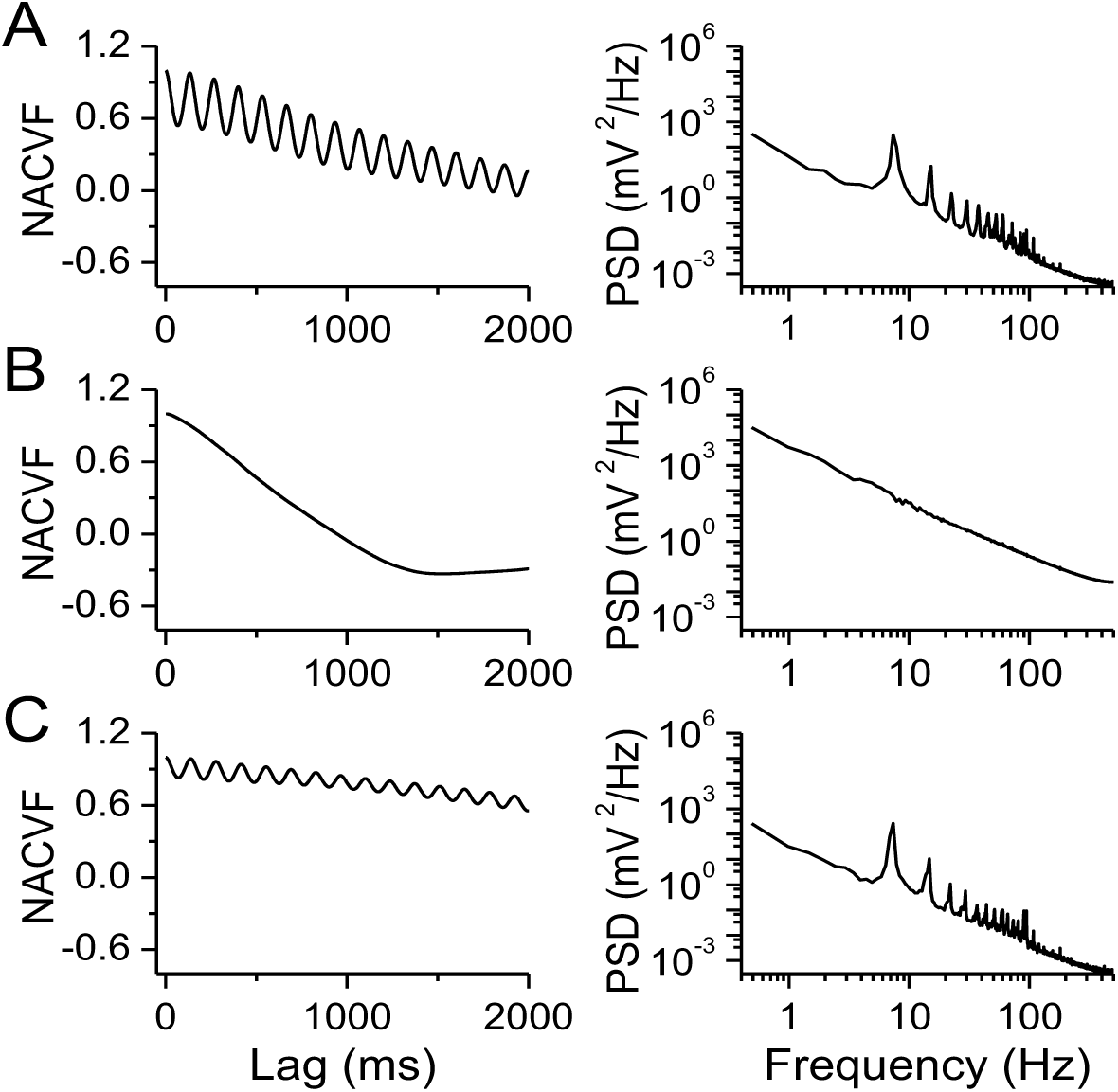
NAI diminishes periodicity of BP. Meca counteracts this effect on BP. Left panels are normalized autocovariance function (NACVF) and right panels are power spectral density (PSD) of BP. (**A**) Pre-nicotine conditions. (**B**) NAI exposure. (**C**) NAI in the presence of Meca.

To quantify the temporal relationship between BP and cardiac activity, we calculated NCCVF of BP vs. ECG. A periodic cross-correlation between BP and ECG was present in pre-nicotine conditions (Fig. 9 A) indicating that instantaneous BP is highly correlated with the heartbeat and that both are rhythmic. During NAI, NCCVF showed a high cross-correlation within one cycle while periodicity rapidly diminished afterward. That NAI disturbed BP periodicity while short latency cross-correlation with cardiac activity persisted suggests that cardiac arrhythmia and irregular cardiac output underlie disruption of rhythmic BP. Following systemic application of Meca, periodicity of BP persisted upon NAI, i.e. Fig 8 C and Fig. 9 C).

**Figure 9.**
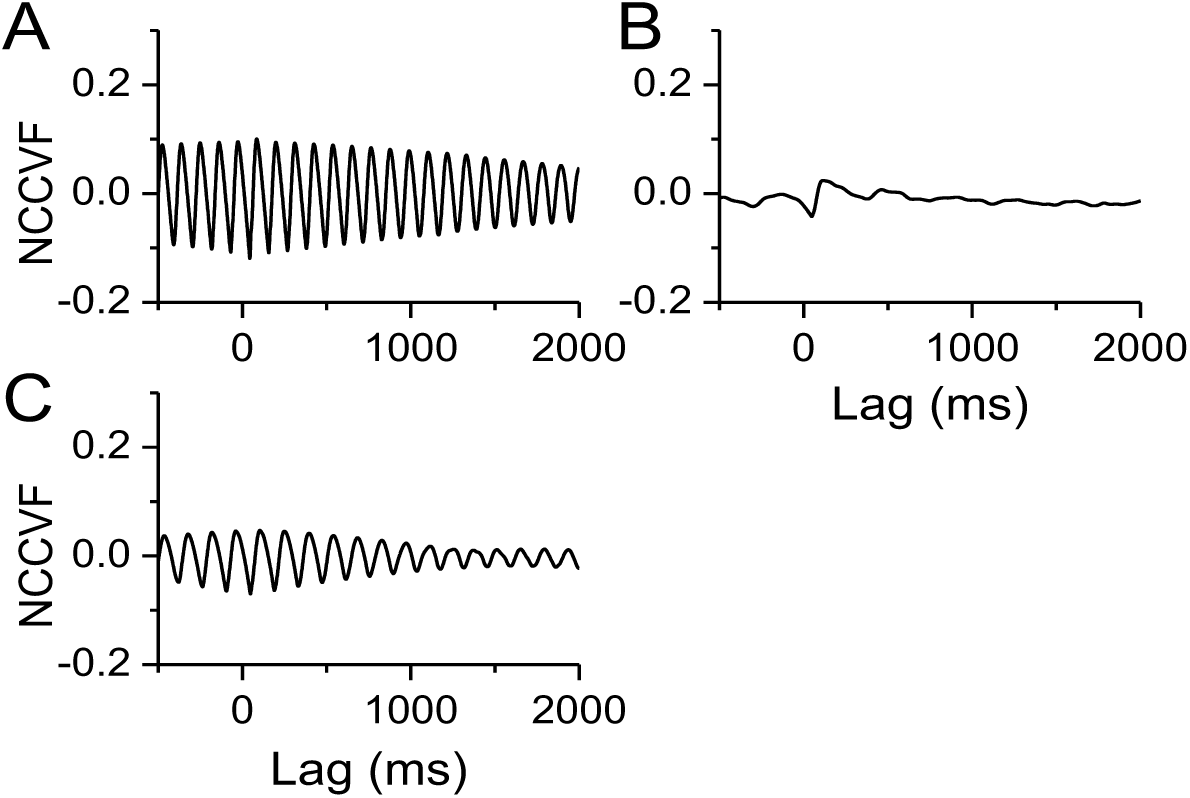
NAI diminishes periodicity of BP while cross-correlation remains within one cycle. Meca counteracts this NAI effect on periodicity. Normalized cross-covariance function (NCCVF) of BP as a continuous process vs. ECG as a point process was calculated. (**A**) Pre-nicotine conditions. (**B**) NAI exposure. (**C**) NAI in the presence of Meca.

## Discussion

Using a newly developed, non-invasive, alveolar region-targeted aerosol inhalation method to episodically deliver nicotine to pregnant rats, we demonstrated that (A) NAI produced PK in both arterial and venous blood in pregnant rats resembling that of smoking a cigarette in humans; (B) NAI induced a transient reduction in mean and diastolic blood flow and in amplitude of pulse flow, as well as irregular fluctuations in diastolic and pulse flow of the uterine artery; (C) Resection of the ovarian nerve counteracts the NAI-induced reduction in mean and diastolic flow in the uterine artery without affecting the high magnitude irregular fluctuation of blood flow and cardiac arrhythmia; (D) NAI induces a transient high magnitude irregular fluctuation of systemic BP, an increase in amplitude and variability of the pulse pressure; (E) NAI induced a transient cardiac arrhythmia including bradycardia and irregular rhythm. ECG abnormalities include a) SA block; b) sinus arrest; c) AV block 2° and 3°; d) ectopic (atrial, AV junctional and ventrical) beats; e) AV junctional or atrial escape rhythm; f) paroxysmal supraventricular tachycardia (SVT); g) ventricular premature beats. (F) nAChR antagonist Meca blocks the nicotine effects on blood flow, BP and cardiac rhythm. Our data suggest that cigarette smoking-relevant NAI stimulates and then desensitizes nAChRs in both sympathetic and parasympathetic systems leading to a striking transient hemodynamic disruption including irregular fluctuation of systemic BP as well as reduction and fluctuation of uterine blood circulation. Episodic uterine hemodynamic disturbances multiple times a day, as would happen in many pregnant smokers due to their smoking patterns, will lead to significant and repeated bouts of uteroplacental and fetal ischemia ^18,19^, which could be a mechanism underlying pregnancy complications and developmental disorders associated with maternal smoking. Furthermore, cardiac arrhythmia and systemic hemodynamic disruption are potential risks for cardiac ischemia, sudden cardiac death and stroke in adult smokers ^2^. These findings indicate substantial health risks of cigarette smoking to cardiovascular function, pregnancy and human development, and challenge the safety of inhalation of pure nicotine as alternatives for cigarette smoking, such as electronic nicotine delivery systems/E-cigarettes and other tobacco products.

Our arterial and venous nicotine PK profiles in rats are consistent with those of Lunell et al ^15^ in non-pregnant adult smokers (male and female) who smoked one cigarette after abstinence for 24 hrs (there has been no similar report on arterial PK in pregnant smokers). We suggest that the rapid rising pattern and the peak values of nicotine concentrations of arterial PK play a major role in the cardiovascular effects we observed, such as arrhythmia, large fluctuations of BP and uterine flow, all novel observations in mammals.

To compare the results obtained in this study with the studies of human smoking on cardiovascular system and hemodynamics, 3 factors should be taken into account. i) cigarette smoke contains > 7000 chemical compounds ^20^; their effects on human systems could be the sum of these components and their interactions. To analyze the effects of the components in cigarette smoke and their interactions is a long-term challenge in the field; ii) the vascular system in human is a much larger buffer system to changes in blood pressure and other hemodynamic parameters; iii) current techniques for measuring human uterine arterial blood flow are limited as mention above. Therefore, to extend our results to human smoking studies await continuous measurement of blood pressure and continuous direct measurement of uterine blood flow during smoking in pregnant women. On the other hand, our results of this study provide clear evidence for the effects of the major active compound nicotine pharmacokinetically equivalent to human smoking on hemodynamics, and also indicate a mechanistic link between maternal smoking and fetal development as well as pregnancy complications.

Our ECG data show that inhalation of nicotine aerosol induces cardiac arrhythmia including SA block, sinus arrests, A-V block 2° and 3°, supraventricular escape rhythm, signs consistent with vagus nerve stimulation. These results are consistent with Hayashi et al ^21^ that nicotine prolonged the interatrial conduction time and effective refractory period that is probably due to the blockade effects of nicotine on the transient outward potassium current (I_to_) ^22^ and on the rapid component of the delayed rectifier current (I_Kr_) ^23^ in atrial myocytes. Nicotine injection (i.v.) in a dose equivalent to 2-fold that of smoking a cigarette in humans induces cardiac arrhythmia in anesthetized dogs ^24^. We found that NAI equivalent to smoking a cigarette in humans induces arrhythmia in awake rats. In rats, the cardiovascular responses, such as heart rate, BP and blood flow, to acute nicotine infusion are not affected by chronic exposure to cigarette smoke^25^. Cardiovascular responses to acute NAI in animals that undergo chronic intermittent inhaled nicotine exposure remain to be determined.

In our previous study, we determined the inhalation toxicity of nicotine LC50 for rats is 2.3 mg/L in air 20 min ^11^. We observed respiratory depression and seizures but those rats were not died of respiratory depression. This current study suggests, in addition to seizures, rats would die of cardiac arrhythmia leading to heart failure in high dose of nicotine inhalation. This study also suggests that the nicotine induced hemodynamic disruption and cardiac arrhythmia in adults can be an important contributor to the epidemiology of increased sudden death rate due to smoking ^2^. Peters and colleagues ^4^ found an association between smoking cessation and reduction in death from cardiac arrhythmia for patients with left ventricular dysfunction after myocardial infarction.

Endogenous ACh or exogenous nicotine activate pre- and postsynaptic nAChRs that facilitates glutamatergic neurotransmission to and depolarizes parasympathetic preganglionic cardiac vagal neurons in the nucleus ambiguus; the postsynaptic effects are mediated by α3β2/α6βX and α3β4 nAChRs ^26,27^. Nicotine also activates postsynaptic α3β2 nAChRs that mediate cholinergic transmission to the parasympathetic ganglionic neurons ^28^ that send vagal efferents innervating the heart. Our results are consistent with these mechanisms of nicotine actions leading to cardiac bradycardia, arrhythmia and fluctuation of arterial BP.

## Methods

### Ethical approval

All animal use procedures were in accordance with The National Institutes of Health (USA) Guide for the Care and Use of Laboratory Animals and were approved by the UCLA Institutional Animal Care and Use Committee. Efforts were made to minimize the number of animals used and their pain. The investigators understand the ethical principles under which *The Journal of Physiology* operates and our work complies with this animal ethics checklist.

### Animals

Pregnant Sprague-Dawley rats were purchased from Charles River Laboratories International, Inc., Wilmington, MA. They were housed in the vivarium under a 12-hr light/dark cycle and had *ad libitum* access to food and water. The pregnant rats of Gestation 16-20 days (E16 – E20) were used for experiments in this study.

### Nicotine aerosol generation and exposure system

A nicotine aerosol generation and exposure system was described in our previous publication ^11^ that included a 3-jet Collison nebulizer (BGI Inc., Waltham, MA), a homemade nose-only exposure chamber and rat holders (CHT-250, CH Technologies Inc. Westwood, NJ). Nicotine aerosols with defined droplet size distribution can be consistently generated in this system. The pressure of the air that enters the nebulizer determines the volume, the droplet size distribution and the concentration of the output aerosol^29^. The air pressure and flow rate were well controlled and continuously monitored during the nicotine aerosol exposure experiments. At the breathing zone of the nose-only exposure chamber, nicotine aerosol droplet size distribution, as described previously ^11^, were determined to be within the range of respirable diameter.Respirable diameter is defined as the aerodynamic diameter of particles capable of reaching the gas exchange region in the lungs (the alveoli) for the organism under study ^14^.

The experiments were carried out in a fume hood at room temperature of 22 ± 1°C. Nicotine (freebase) was dissolved in 0.7% NaCl solution for an osmolality ∼300 mOsm/kg. pH was adjusted with HCl to pH 8.0. Air pressure for aerosol generation was 20 psi.

### Arterial and venous blood sample collection and plasma nicotine assay

For arterial blood sample collection, femoral artery pre-catheterized pregnant rats (E17) were purchased from Charles River Inc. For venous blood collection, local anesthetic cream was applied to the rat tail 30 min prior to insertion of the catheter. The rat was put into the rat holder, and the tail was warmed with warm water or an infrared light. A temporary catheter (24G) was inserted into the lateral tail vein. Warm saline (2.5 mL) was infused into the arterial or venous catheter for blood volume compensation before blood collection. Then 0.1-0.2 mL of Heparin (100U/mL) was infused.

The 1^st^ sample as pre-nicotine control was collected. Then nicotine aerosol was generated from the nebulizer and at the same time, a timer was started. The rat was exposed to the nicotine aerosol for 2 min. Blood samples (equivalent to 0.2 mL plasma) were collected at a series of time points (see Results) with 0.6 mL capillary blood collection tubes containing EDTA-K2 (RAM scientific Inc. Yonkers, NY). Venous bleeding from the catheter can be greatly facilitated by massaging the tail over the vein.

Samples were centrifuged within 15 min of collection at 1000 g (gravity) for 12 min at room temperature. Supernatant plasma was transferred to labeled Eppendorf tubes and store below -20°C. The plasma samples were sent to NMS Labs (Willow Grove, PA) where nicotine and its metabolite cotinine were measured with High Performance Liquid Chromatography/Tandem Mass Spectrometry (LC-MS/MS).

### Measurement of Uterine artery blood flow

Uterine artery blood flow was measured with a chronically implanted perivascular flowprobe using transit-time ultrasound technology (Model 0.5PSB, Transonic Systems Inc. NY, USA) ^12,30^.

Pregnant rats (E16 - E17) were anesthetized with isoflurane (3.5% induction and 1.5-2.0% maintenance). A 1.5 cm incision on the mid-scapular area was made to produce the exit of the micro flowprobe connector. Then, a blunt dissection of subcutaneous tissue to the abdominal region was performed to create a subcutaneous tunnel. The rat was placed in a dorsal recumbency, and a midline abdominal skin incision (∼ 3 cm) was made. An incision through the linea alba was made next. The intestines were deflected to the left side of the rat to expose the uterine horns. The right uterine horn was exposed. During the whole procedure the uterine horn was irrigated with warm sterile saline solution. A segment of ∼ 2 mm of the uterine artery, between two segmental arteries rostral to the uterine cervix, was separated free from mesometrium and uterine vein. The uterine artery was gently lifted and placed in the lumen of the flowprobe. The space between the probe and uterine artery was filled with an acoustical couplant gel to ensure a good recording of the blood flow signal. The flowprobe cable was sutured (Vicryl, 5-0) on the uterine horn to stabilize the probe. The flowprobe connector at the other end of the cable was gently grasped and was guided to the mid-scapular incision trough the subcutaneous tunnel. The connector cuff was sutured to the skin, and the incisions were sutured close with silk (4-0). The abdominal wall was closed with continuous absorbable suture (Vicryl, 5-0). At the end of the experiment, antibiotic (Animax cream, Pharmaderm) was applied over the sutures, and the analgesic Carprofen (5 mg/kg) was applied subcutaneously. Animals were allowed to recover in a warmed environment.

The perivascular flow probe was connected to a flow meter (TS420, Transonic Systems Inc. Ithaca, NY, USA) where the flow signal was low-pass filtered at 40 Hz and the scale factor was 1.5 ml/min/V or 6 ml/min/V.

### Ovarian nerve resection

In the rat, the ovarian nerve is in parallel with the ovarian vessels ^17^. In the region of the anastomosis between the ovarian and uterine vessels, the ovarian nerve turns to follow the uterine artery and vein. In vivo, the ovarian nerve is too small and transparent (30-40 μm in diameter) to be seen without staining. To completely disconnect the nerve, we carefully eliminated all the tissue around both ovarian vessels under a dissection microscope. Approximately a segment of 4 mm of the ovarian artery and vein were dissected free of surrounding tissue at approximately 1 cm from the anastomosis between the ovarian and uterine arteries.

After the surgery for implanting the flowprobe and cutting the ovarian nerve of the same side, animals were allowed to recover for two days before experiments. Uterine artery blood flow and ECG were recorded from awake, restrained rats (in a rat holder) in NAI experiments (Rats were E18-E20 on experiment days).

### Arterial blood pressure and ECG recordings

Unilateral femoral artery pre-catheterized pregnant rats were purchased from Charles River Laboratory (Charles River Laboratories, Inc.). The artery catheter (volume ∼ 60 μl) was filled with saline containing heparin (50-100 IU/ml) and was connected to a pressure transducer (NL 108T2, Neurolog^TM^ System, Digitimer Limited, Letchworth Garden City, UK) with tubing filled with saline. The transducer was connected to a pressure amplifier (NL 108A, Digitimer Limited) with a gain of 1V/100mmHg.

ECG was recorded with two subcutaneous needle electrodes placed on the left front and rear limbs (lead III). ECG signal was amplified (x10,000) and band-pass filtered at 1-300 Hz with an AC amplifier (Model LP511, Grass/Natus Neurology Inc. Warwick, RI, USA).

Blood flow, BP and ECG signals were acquired at a sampling frequency of 1K Hz using a digitizer MiniDigi 1A and software Axoscope 10.2 (Axon/Molecular Devices, LLC. Sunnyvale, CA. USA).

### Chemicals

Nicotine used in this study was (s)-(-)-nicotine freebase (liquid, 99%) purchased from Alfa Aesar Co, Ward Hill, MA. Mecamylamine-HCl was from Tocris Bioscience, Bristol, UK.

### Data analysis

Since blood flow and BP were pulsing, we analyzed blood flow, BP and ECG by summarizing parameters of continuous records at different conditions (40 sec for pre- and post-aerosol delivery and 30 sec during nicotine aerosol delivery). Summary parameters (expressed as mean ± SD of n animals in the text and mean ± SE in figures) were compared in two-way repeated measures ANOVA followed by post hoc multiple comparison method of Sidak using the software SAS (v 9.4, SAS institute Inc. Cary, NC, USA)

To quantify the regularity of the pulsing processes of blood flow, BP and ECG, coefficient of variation (CV) and normalized autocovariance function (NACVF), power spectrum and normalized cross-covariance function (NCCVF) ^31^ were calculated.

CV = SD/mean

For digital signal analysis, we consider discrete-time series. With the assumption of stationarity and ergodicity, autocovariance function of a time series Y(m):

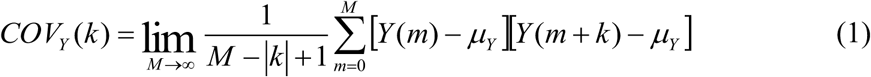

m = 0, 1,…M. k =…-1, 0, 1,…. μ^Y^ is the mean of Y(m). NACVF is

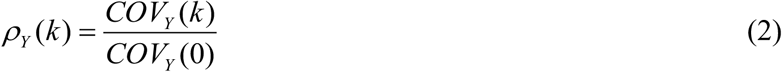

COV_Y_(0) equals the variance of Y(m).

To quantify the correlation between uterine artery blood flow or BP and ECG, we take QRS waves as point events and therefore, an ECG recording as a point process. We have:

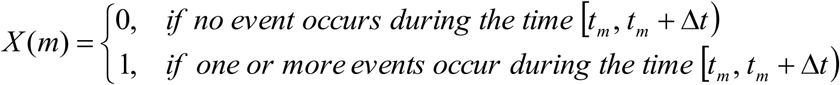

In practice, Δt is taken so small that the probability of more than one events occurring during [t_m_, t_m_ + Δt) is zero. The cross-covariance function of a continuous process (blood flow or BP) Y(m) vs. a point process X(m) (both X(m) and Y(m) are discrete-time series) is defined as:

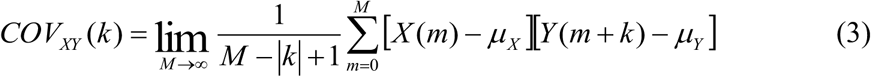

For a point process, μX is the mean frequency of points. NCCVF is

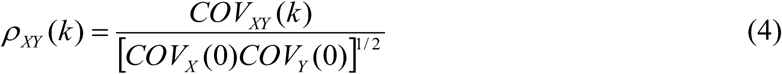

For a point process,

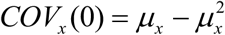

NACVF and NCCVF represent strength of correlation as a function of time on a monotonic scale ranging from -1 to +1 ^31^. Software Clampfit 10 (Molecular devices) and DataView 9 (W. J. Heitler, University of St. Andrews, St. Andrews, UK) were used for these analyses.

## Acknowledgments

This work was supported by California Tobacco-Related Disease Research Program Grant (18XT-0183) and National Institute of Health (NIH) Grants (1R43DA031578, R44 DA031578-02 and R01HL040959). We thank Dr. Qiong Li for her advice and assistance with the ECG data analysis.

## Author contributions

X.M.S. and J.L.F. conceived and designed experiments. X.M.S., H.E.L-V. and J.L. performed the experiments and analyses. X.M.S., H.E.L-V. and J.L.F. interpreted the results and wrote the manuscript. All authors have approved the final version of the manuscript and agreed to be accountable for all aspects of the work.

